# Stabilization Mechanism of Initiator Transfer RNA in the Small Ribosomal Subunit from Coarse-Grained Molecular Simulations

**DOI:** 10.1101/2024.07.12.603193

**Authors:** Yoshiharu Mori, Shigenori Tanaka

## Abstract

Proteins play a variety of roles in biological phenomena in cells. Proteins are synthesized by a ribosome, which is a large molecular complex composed of proteins and nucleic acids. Among many molecules involved in the process of protein synthesis, transfer RNA (tRNA) is one of the essential molecules. In this study, coarse-grained molecular dynamics simulations were performed to understand how the tRNA molecule is stabilized in the ribosome, and the free energy along the dissociation path of the tRNA was calculated. We found that some ribosomal proteins, which are components of the ribosome, are involved in the stabilization of the tRNA. The positively charged amino acid residues in the C-terminal region of the ribosomal proteins are particularly important for the stabilization. These findings contribute to our understanding of the molecular evolution of protein synthesis in terms of the ribosome, which is a universal component of life.

**TOC Graphic:** 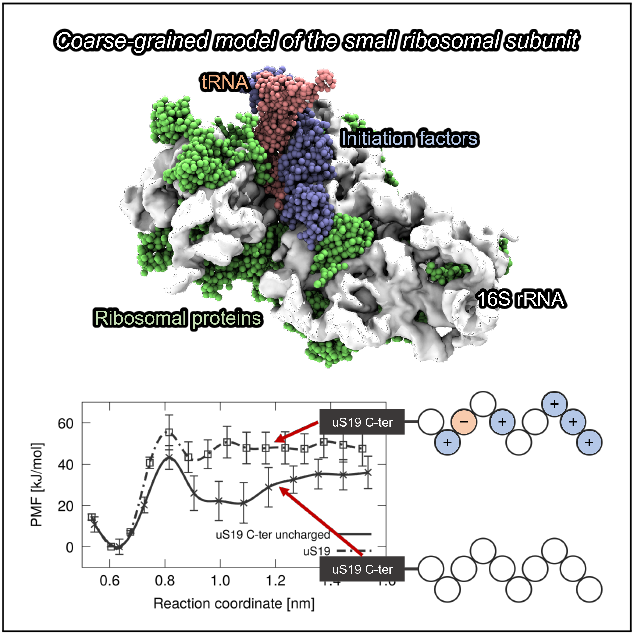

---

Ribosomes are large complexes of protein and RNA molecules that are responsible for protein synthesis in cells (Fig. 1a). The genetic information in DNA is transcribed into messenger RNA (mRNA). Proteins with sequences corresponding to the mRNA are synthesized in the ribosome. The process of synthesizing proteins from such RNA is called translation. The process of protein synthesis in the ribosome is mainly divided into initiation, elongation, and termination.^1,2^ While the elongation process is the actual process of protein synthesis in which peptide bonds are formed,^3^ the initiation and termination processes are also essential for high-quality protein synthesis.

**Figure 1:**
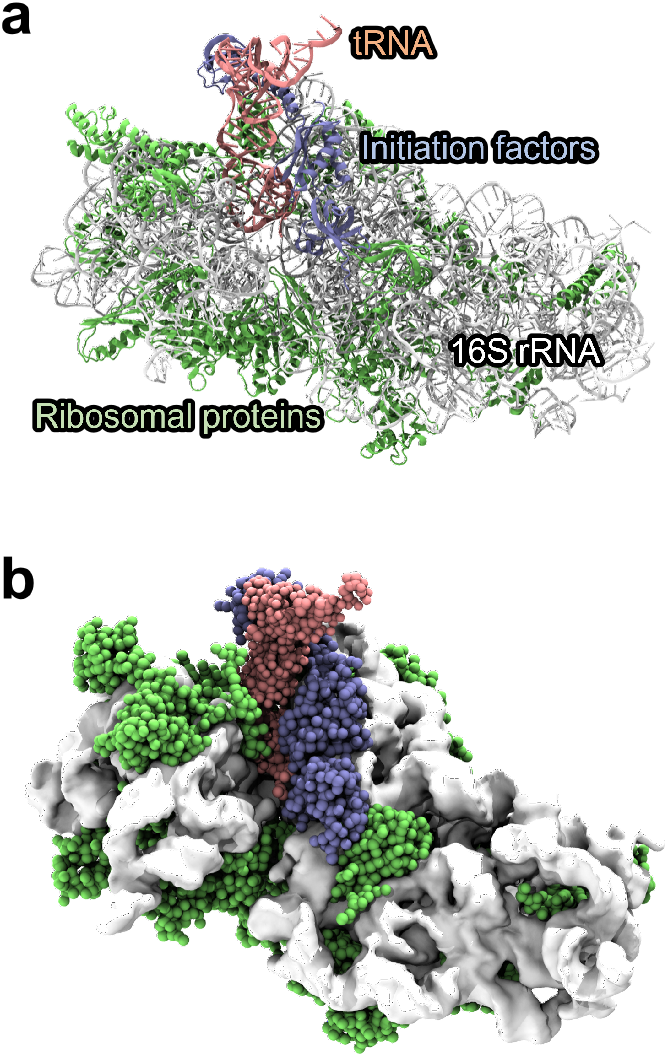
Three-dimensional structure of the PIC (preinitiation complex). These figures are rendered by VMD.^4^ (a) The structure of the PIC is shown as a cartoon representation. 16S rRNA, tRNA, ribosomal proteins, and initiation factors are shown in white, pink, green, and blue, respectively. (b) A coarse-grained model of the PIC is shown. The ribosomal RNA (16S rRNA) of the PIC is shown as a surface representation.

In this study, we focused on the ribosome in the initiation process in bacteria.^5,6^ The initiation process serves as a check process for the subsequent elongation process. In the initiation process, a preinitiation complex (PIC) is formed. The PIC consists of the small ribosomal subunit, which is composed of ribosomal RNA and ribosomal proteins, initiation factors (IFs), and initiator transfer RNA (tRNA). The initiation factors are proteins that are specific for initiation. In the case of bacteria, there are IF1, IF2, and IF3. Experimental structure analyses suggest that some ribosomal proteins are involved in stabilizing the initiator tRNA in the ribosome,^7,8^ although the terminal region of these proteins does not appear to have secondary structure.

The association of the initiator tRNA with the ribosome is necessary for the initiation of protein synthesis. Because there are many different types of tRNA in the cell, it is necessary for the PIC to be able to selectively bind to the initiator tRNA. In other words, the initiator tRNA should be stably present in the PIC. The purpose of this study is to understand how the tRNA is stabilized in the PIC. Each translation process in the ribosome is known to be evolutionarily conserved in bacteria,^5^ archaea,^9^ and eukaryotes.^10^ The corresponding initiation factors and ribosomal proteins exist for the three domains. Insights into the mechanism of tRNA stabilization in the ribosome lead to an understanding of the evolution of the molecular mechanisms of protein synthesis. ^11^

Since proteins play a major role in cellular functions, translation is an important process in cells. To understand the translation in detail, the three-dimensional structures of ribosomes have been elucidated.^3,7,12–16^ Such studies have revealed, for example, that RNA functions as a catalyst in the formation of peptide bonds.^3^ The structure of the PIC has also been determined to understand the translation initiation process. ^7,13,15^ Such structural studies have suggested that the initiator tRNA in the ribosome interacts with many of the molecules that form the PIC. In addition, molecular simulations have been used to understand biomolecular systems in detail. However, ribosomes are much larger than the biomolecules that are typically studied using molecular simulations. Therefore, structure-based or coarse-grained models are often used to treat such systems.^17,18^

In this study, we performed molecular simulations using a coarse-grained model including the complete PIC system to clarify which molecules and interactions are responsible for stabilizing tRNA in the PIC. The results of the simulations provided a novel insight into the free energy landscape of the initiator tRNA within the ribosome. In addition, it has recently become possible to obtain the structure of large biomolecular assemblies not only by experiment but also by computational modeling.^19,20^ The coarse-grained simulations in this study demonstrate that even large biomolecular assemblies obtained with these methods can be analyzed to study detailed molecular interactions.

Because the PIC is a molecular complex consisting of more than 20 protein and RNA molecules, it is difficult to perform all-atom molecular simulations as described above. Therefore, we performed molecular dynamics (MD) simulations using a coarse-grained model of the protein-RNA complex. The following is a summary of the simulation protocols (for details, see Computational Details in Supporting Information). The PIC in bacteria was used as the simulation system, and the initial structure of the complex was the three-dimensional structure obtained by cryo-electron microscopy.^7^ The PIC consisted of the small ribosomal subunit, mRNA, initiator tRNA, and the initiation factors IF1 and IF3. To model amino acid residues not determined by the experiments, we performed structure modeling using MODELLER^21^ and Chimera.^22^ We used the Martini 2.2 force field for proteins,^23–26^ RNAs,^27^ and solvent. The coarse-grained PIC using the Martini model consisted of approximately 16,000 beads (Fig. 1b). We added coarse-grained beads corresponding to the solvent and ions to the coarse-grained PIC system. The total number of beads in the simulation system was about 170,000. The structural analyses of the PIC suggested that the PIC has several structural states during the tRNA binding process. ^7^ To compare the different states that PIC can take, three states were considered in this study. Following the reference of the experiment,^7^ we refer to these states as PIC-2A, PIC-3, and PIC-4 in this letter. PIC-2A is a state that occurs immediately after the initiator tRNA is bound to the small ribosomal subunit (pdb id: 5lmq). PIC-3 is an intermediate state that occurs after the initiator tRNA is bound to the small ribosomal subunit (pdb id: 5lmt). PIC-4 is a state in which the initiator tRNA adapts to the environment of the ribosome (pdb id: 5lmu). We generated a coarse-grained model of the PIC for each of these states as described above.

We performed structural optimization and MD simulations for these simulation systems using GROMACS^28^ and PLUMED.^29,30^ In these calculations, an elastic network model was used to restrain the protein and RNA structures to the experimental structure.^31^ After optimizing the coarse-grained PIC model, an MD simulation for equilibration was performed at 310 K and 0.1 MPa using the temperature and pressure control algorithms.^32,33^ The time step for the time evolution was set to 20 fs. To understand how the components of the PIC contribute to the stability of the initiator tRNA, we calculated the dissociation free energy of the initiator tRNA using the following procedure. The reaction coordinate characterizing the dissociation process of the initiator tRNA was defined by the distance between the tRNA and the small ribosomal subunit (Fig. 2a). First, we defined the set of tRNA beads that are within 0.8 nm of the ribosomal RNA in the initial structure. Similarly, we defined the set of ribosomal RNA beads that are within 0.8 nm of the initiator tRNA in the initial structure. We then calculated the center of mass for the two groups and defined the reaction coordinate as the distance between the two centers. To estimate the dissociation path of the tRNA from the small ribosomal subunit, we performed several 500 ns metadynamics simulations using the reaction coordinate defined above and an additional reaction coordinate. While metadynamics is an effective method for performing efficient structural sampling,^34,35^ it is necessary to carefully design the protocol to accurately calculate the free energy.^35^ Umbrella sampling^36^ is a useful method for accurate free energy calculation. However, we need to define appropriate reaction coordinates that represent the reaction path. In this study, we performed free energy calculations using these two methods to accurately estimate the free energy along the reaction coordinate.^37–39^ Umbrella sampling simulations were performed for the reaction coordinate in the range of 0.5 nm to 1.5 nm along the tRNA dissociation path estimated by metadynamics. The total simulation time of the umbrella sampling simulations in PIC-2A, PIC-3, and PIC-4 was 63.0 *µ*s, 84.0 *µ*s, and 57.0 *µ*s, respectively. We used the multistate Bennett acceptance ratio method^40^ to estimate the free energy along the reaction coordinate from the results of the umbrella sampling simulations. We also refer to the free energy for this reaction coordinate as the potential of mean force (PMF).

**Figure 2:**
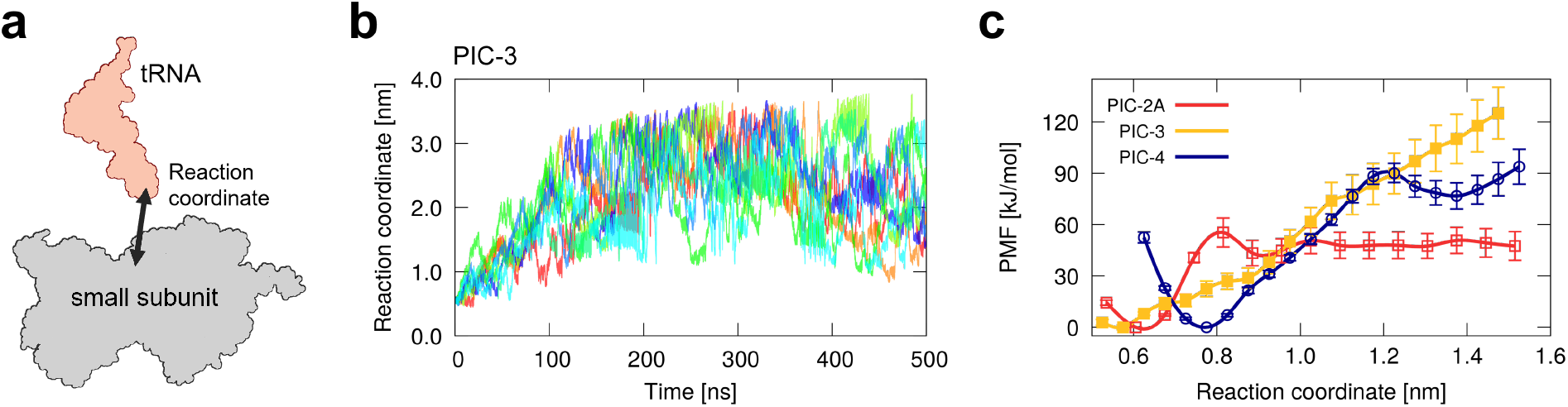
Results of the metadynamics and umbrella sampling simulations. (a) The reaction coordinate is shown schematically. The reaction coordinate is defined by the distance between the tRNA and the small ribosomal subunit (see main text). (b) The time series of the reaction coordinate in the metadynamics simulations in PIC-3 are shown. The eight trajectories are displayed in different colors. (c) The free energy (or PMF) of tRNA dissociation along the reaction coordinate is shown for PIC-2A, PIC-3, and PIC-4. The error bars represent the standard error of the free energy.

Metadynamics simulations were performed to estimate the dissociation path of the initiator tRNA. Figure 2b shows the trajectories of the reaction coordinate in PIC-3. The eight trajectories of the reaction coordinate are shown for the process of tRNA dissociation from the binding region. Although the trajectories are different in each metadynamics simulation, the initiator tRNA was able to dissociate from the binding region within the simulation time in all simulations. Similar results were obtained for PIC-2A and PIC-4 (see Fig. S1 in Supporting Information).

Umbrella sampling simulations were performed using the tRNA dissociation path obtained from the metadynamics simulations, and the free energy along the dissociation path was estimated from these results. Figure 2c shows the free energy along the tRNA dissociation path. It can be seen that in PIC-2A there was a stable tRNA bound state (at about 0.6 nm). In addition, there was a free energy barrier (at about 0.8 nm) in the direction of tRNA dissociation. In PIC-3, in contrast to PIC-2A, it is difficult to identify a clear stable position for the tRNA. In addition, the free energy difference between the tRNA bound state in the ribosome and the dissociated state was larger than for PIC-2A. This suggests that once the initiator tRNA is correctly bound to the small ribosomal subunit, the structure of the PIC in PIC-3 prevents the tRNA from dissociating. In PIC-4, there was a stable tRNA bound state (at about 0.8 nm) similar to PIC-2A. There was also a free energy barrier (at about 1.2 nm) that was higher than that of PIC-2A. In addition, the free energy difference between the tRNA bound state and the tRNA dissociated state was larger than that for PIC-2A. These results indicate that the initiator tRNA can be stably located in the small ribosomal subunit in PIC-4 and is difficult to dissociate from the binding region.

Next, to identify the molecules that contribute to the stabilization of tRNA, we analyzed the distances between the initiator tRNA and nearby molecules along the reaction coordinate. We used MDAnalysis^41,42^ for these analyses. Figure 3a shows proteins near the initiator tRNA, and Figs. 3b-d plot the distances between the tRNA and the proteins along the reaction coordinate in PIC-2A, PIC-3, and PIC-4. uS9, uS13, and uS19 are the names of ribosomal proteins.^43^ Figure 3e shows RNA residues near the initiator tRNA, and Figs. 3f-h plot the distances between the tRNA and the RNA residues in PIC-2A, PIC-3, and PIC-4 along the reaction coordinate. The distance between the tRNA and a molecule is defined as the minimum distance between the beads of the tRNA and those of the molecule. The tRNA formed a codon-anticodon pair with the mRNA along the reaction coordinate in all states (Figs. 3f-h). Similarly, the attractive interaction between the tRNA and the ribosomal protein uS13, and that between the tRNA and the initiation factor IF3 were always present (Figs. 3b-d). In PIC-2A, uS9 interacted with the tRNA in the tRNA binding region (Fig. 3b), whereas in PIC-3, the interaction was reduced upon dissociation (Fig. 3c). In PIC-4, uS9 maintained an interaction with the tRNA (Fig. 3d). This behavior is consistent with the free energy landscape for each state. Therefore, we suggest that the interaction between the tRNA and uS9 is important for stabilizing the tRNA. uS19 did not interact with the tRNA in PIC-2A (Fig. 3b). However, in PIC-3 and PIC-4, there was an interaction between the tRNA and uS19 (Figs. 3c and 3d). One of the reasons why the tRNA in PIC-4 was more stable than that in PIC-2A is the interaction between the tRNA and uS19. We then examined the interactions between the initiator tRNA and the RNA residues. Interactions between the initiator tRNA and several ribosomal RNA residues are conserved in bacteria,^7,44^ archaea,^45^ and eukaryotes.^46^ We found that G1338 and A1339 had interactions with the tRNA in the tRNA binding region and that the distance was longer in PIC-4 than in PIC-2A (Figs. 3f and 3h). While A790 did not significantly interact with the tRNA in PIC-2A (Fig. 3f), the intermolecular distance in PIC-3 and PIC-4 was shorter than that in PIC-2A (Figs. 3g and 3h), showing that an attractive interaction between the tRNA and A790 was formed. Although the distance for C1400 shows a similar tendency to that for G1338 and A1339, the interaction with tRNA was maintained in PIC-3 and PIC-4 (Figs. 3g and 3h) in contrast to PIC-2A (Fig. 3f).

**Figure 3:**
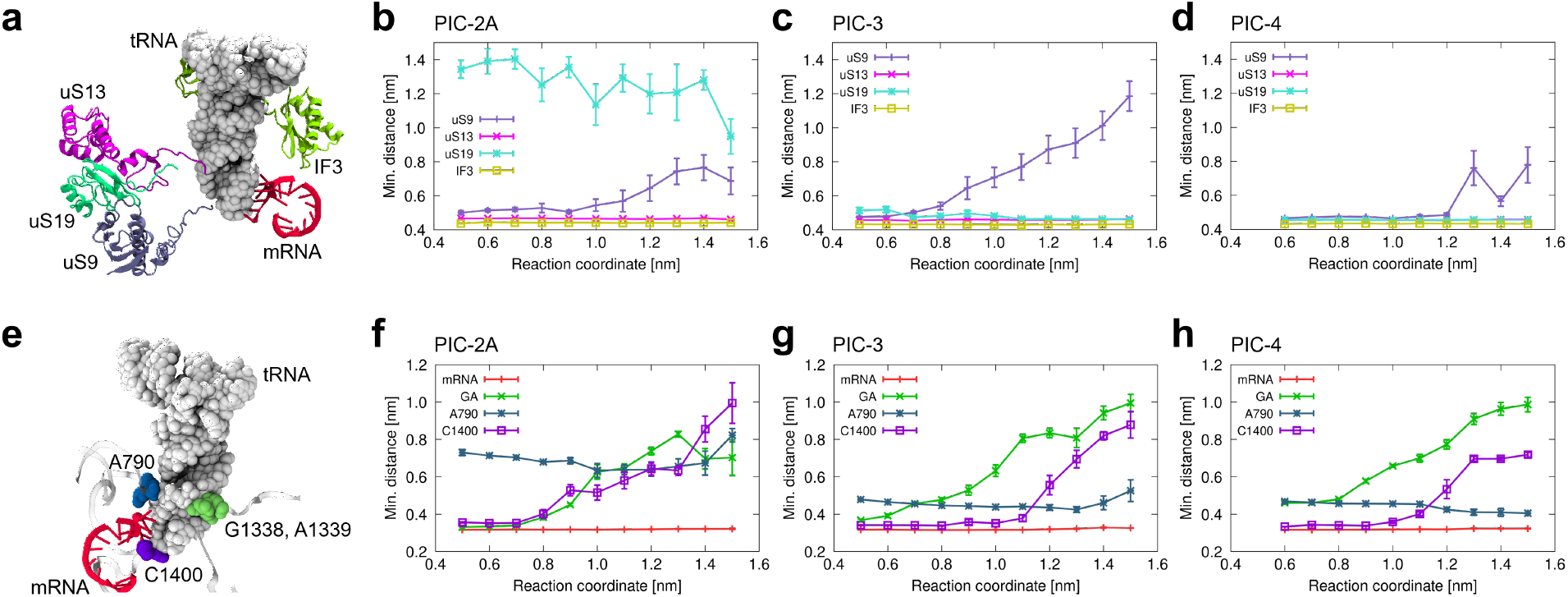
Interactions between the tRNA and the surrounding molecules. (a) The ribosomal proteins near the tRNA are shown. The distances between the tRNA and the ribosomal proteins are shown as a function of the reaction coordinate for (b) PIC-2A, (c) PIC-3, and (d) PIC-4. For the definition of the distance, see main text. (e) The ribosomal RNA residues near the tRNA are shown. The distances between the tRNA and the mRNA and ribosomal RNA residues are shown as a function of the reaction coordinate for (f) PIC-2A, (g) PIC-3, and (h) PIC-4. The error bars represent the standard error of the distance.

The above analyses show that the initiator tRNA is stable in the binding region in PIC-2A and PIC-4. Because some ribosomal proteins are located near the tRNA, we consider the effect of the ribosomal proteins on the stability of the tRNA in more detail. Ribosomal proteins form a network structure within the ribosome.^8,47,48^ The terminal region of several ribosomal proteins lacks secondary structure. However, structural analyses suggest that their terminal regions interact with tRNA. In addition, bioinformatics analyses revealed that ribosomal proteins have characteristic amino acid compositions and that some of the proteins contain intrinsically disordered regions. ^49^ Ribosomal proteins are rich in positively charged amino acids such as arginine and lysine. Thus, there can be an attractive interaction between ribosomal proteins and tRNA, which is rich in negative charge. In one experiment, truncation of the C-terminal region of the ribosomal protein uS19 had a negative effect on cell growth.^50^ This suggests that removal of the C-terminal region of uS19 leads to a loss of ribosome function, which could be caused by reduced interactions between the initiator tRNA and uS19. Therefore, the positively charged amino acids in the C-terminal region of ribosomal proteins can contribute to the stability of the initiator tRNA.

To understand how the charged amino acids in the C-terminal region of ribosomal proteins affect the stability of the initiator tRNA, we performed MD simulations for PIC-2A and PIC-4 in which the electrostatic interactions caused by the C-terminal region of uS19 were eliminated. The simulation procedure is summarized as follows. The ribosomal protein focused on in this study was uS19 (see Fig. 3a). To eliminate the electrostatic interactions in the C-terminal region, the electric charges of the last 10 amino acid residues (GKEAKATKKK) in the C-terminal region were set to zero (Fig. 4a). Under these conditions, metadynamics and umbrella sampling simulations were performed to calculate the dissociation free energy of the initiator tRNA. We used the same simulation protocols as described above for the metadynamics and umbrella sampling simulations.

**Figure 4:**
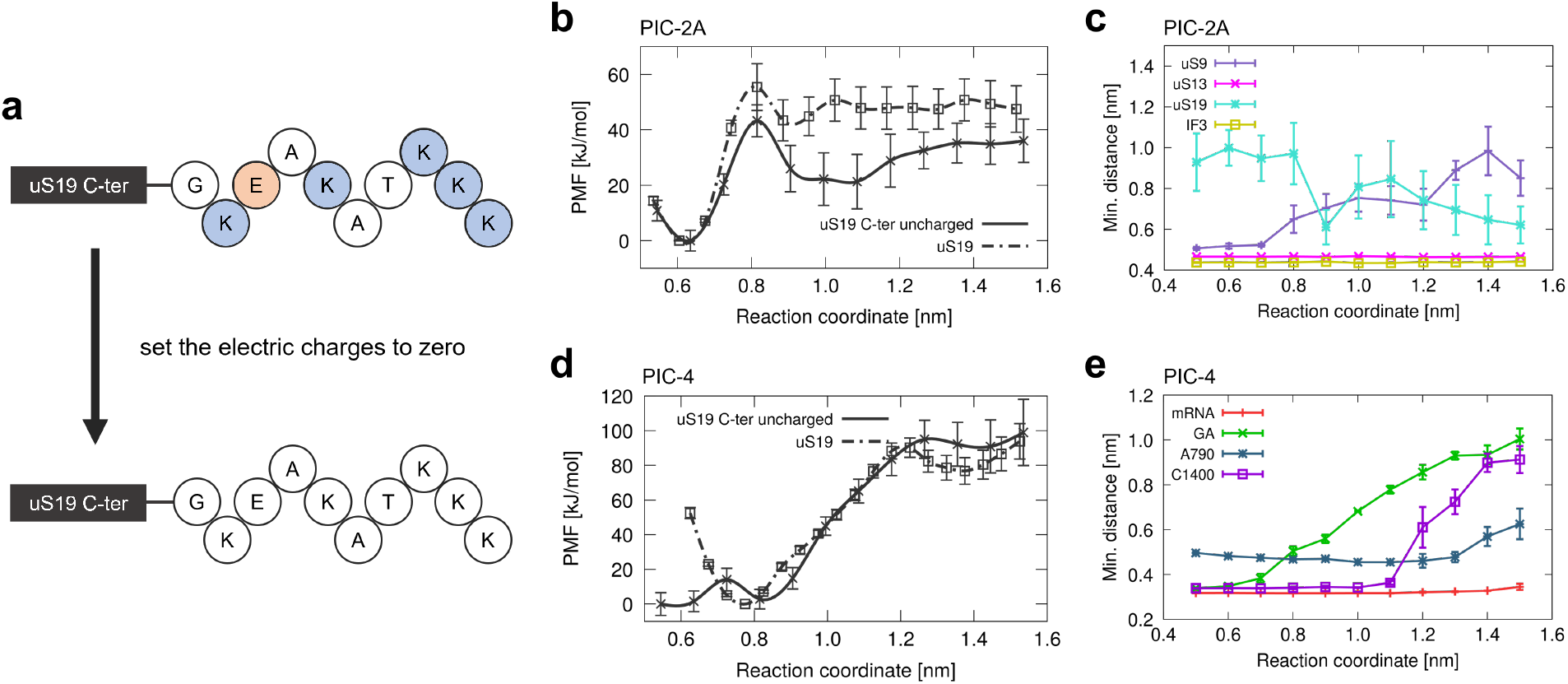
Results of the MD simulations in which the electric charges of the C-terminal residues of uS19 were set to zero. (a) The charge elimination of the C-terminal residues of uS19 is shown schematically. Circles filled with blue, red, and nocolor represent an amino acid residue whose charge state is positive, negative, and neutral, respectively. Each alphabetic character represents the name of an amino acid residue. For (b) PIC-2A and (d) PIC-4, the free energy (or PMF) of tRNA dissociation along the reaction coordinate is shown for the two charge states. The error bars represent the standard error of the free energy. For (c) PIC-2A and (e) PIC-4, the distances between the tRNA and the surrounding molecules are shown as a function of the reaction coordinate. The error bars represent the standard error of the distance.

The results of the free energy calculations for PIC-2A are shown in Fig. 4b. We can see that the binding region of the initiator tRNA in the ribosome was the same whether the C-terminal region of uS19 had the electric charges or not. On the other hand, the height of the free energy barrier of tRNA dissociation was different for each condition. When the charges in the C-terminal region of uS19 were absent, the barrier height was lower than when the charges in the C-terminal region were present. The same holds for the free energy difference between the bound and dissociated states of the tRNA. We found that the electrostatic interaction between the initiator tRNA and the C-terminal region of uS19 contributes to the increase in the free energy barrier. This means that the bound tRNA is easily dissociated from the ribosome when the charges in the C-terminal region of uS19 are absent. Figure 4c plots the distances between the tRNA and the proteins in PIC-2A along the reaction coordinate. Compared to the case where the C-terminal region of uS19 was charged (see Fig. 3b), uS19 was closer to the tRNA. This result seems to be inconsistent with the removal of electrostatic interactions with the tRNA. This could be because the repulsive interactions between uS19 and other ribosomal proteins were also eliminated. We also found that uS9 was more separated from the tRNA around the free energy barrier (Fig. 4c), which is a different result from that obtained when the C-terminal region of uS19 was charged (Fig. 3b).

The results of the free energy calculations for PIC-4 are shown in Fig. 4d. Compared with PIC-2A, the free energy difference between the bound and dissociated states of the initiator tRNA was not significantly affected by the absence of charges in the C-terminal region of uS19. This suggests that in PIC-4, interactions between the tRNA and the C-terminal region are less important during dissociation. We also found that the free energy landscape in the tRNA binding region differed between the two charge states. When the C-terminal region was charged, the tRNA was stable with its position displaced. However, when the C-terminal region was not charged, the tRNA did not exhibit a well-defined bound state in the ribosome, suggesting that the tRNA is unstable within the ribosome. In PIC-4, the electrostatic interactions between the initiator tRNA and the C-terminal region of uS19 are necessary to place the tRNA in its correct position within the ribosome. Figure 4e plots the distances between the tRNA and the RNA residues in PIC-4 along the reaction coordinate. There is not significant difference compared to the results obtained when the C-terminal region of uS19 was charged (see Fig. 3h). The main difference is that the distances between the tRNA and the RNA residues (G1338 and A1339) were shorter than those when the C-terminal region was uncharged. This difference explains why the tRNA could not be located in the correct position within the ribosome when the C-terminal region was uncharged.

We have analyzed and discussed the effects of the charge state in the C-terminal regions of uS19 for PIC-2A and PIC-4. As shown in Figs. 4b and 4d, even the electrostatic interactions of the C-terminal region of a single ribosomal protein have a significant effect on the stabilization of tRNA. In the case of PIC, the three-dimensional structure shows that at least uS9 and uS13, in addition to uS19, interact with the tRNA through their C-terminal regions. This suggests that ribosomal proteins play an important role in the stabilization of the initiator tRNA in the ribosome, as well as in the association and dissociation processes of the tRNA.

In this study, we performed molecular dynamics simulations using a coarse-grained model to understand the stabilization mechanism of the initiator tRNA when the translation preinitiation complex is formed. The ribosome needs to form a proper structure for the translation process to occur correctly. It is important that the initiator tRNA is bound to the correct location within the ribosome. From the results of the free energy analyses along the tRNA dissociation path, we found that the tRNA can be stable within the ribosome and that the electrostatic interactions between the tRNA and several ribosomal proteins are essential. This fact is supported by the results of MD simulations performed to understand the effect of charges in the C-terminal region of uS19. To summarize, the interactions with mRNA, ribosomal RNA, and ribosomal proteins close to the tRNA contribute to the stabilization of the initiator tRNAs in the ribosome. Insight into the various interactions between tRNAs and ribosome components will lead to an understanding of the molecular mechanism of accurate protein synthesis during the translation process. These findings will also be useful in understanding the mechanism of selection of cognate tRNAs from the many types of tRNAs in cells during the elongation process, as well as the initiation process. By identifying conserved interactions within the ribosome of bacteria, archaea and eukaryotes, we expect to understand the origin of life through translation in the ribosome. ^51^

## Acknowledgement

We acknowledge the Grants-in-Aid for Scientific Research from the Ministry of Education, Culture, Sports, Science and Technology (MEXT) (No. JP17H06353) and Japan Society for the Promotion of Science (JSPS) (Nos. JP18K03825, JP21K06098, JP21K06113), and MEXT Quantum Leap Flagship Program (No. JPMXS0120330644). The computation was partly performed using Research Center for Computational Science, Okazaki, Japan (Project: 20-IMS-C261, 21-IMS-C013, 22-IMS-C014).

## Supporting Information Available

The following file is available.

- Supporting_Information.pdf: Computational details and additional results of the simulations.

## Notes

### Competing Interest Statement

The authors have declared no competing interest.

