## Supporting Information for "Stabilization Mechanism of Initiator Transfer RNA in the Small Ribosomal Subunit from Coarse-Grained Molecular Simulations"

### 1. Computational Details

#### 1.1 System preparation

1.1.1 *Molecular complex for simulations: translation preinitiation complex*

We used a translation preinitiation complex (PIC) from *Thermus thermophilus* as the molecular complex for simulations. The initial structure of the complex was the three-dimensional structure obtained by cryo-electron microscopy [1]. This experiment determined eleven structures of the PIC, which represents distinct steps during initiation. We chose three structures corresponding to different steps from the eleven structures. These structural states were PIC-2A, PIC-3, and PIC-4, where we refer to the names of the states as the same as in the experimental study [1]. The PDB IDs of PIC-2A, PIC-3, and PIC-4 are 5lmq, 5lmt, and 5lmu, respectively.

The PIC consisted of the small ribosomal subunit, mRNA, initiator tRNA, and the initiation factors IF1 and IF3 (except for PIC-4, see Table S1). The small ribosomal subunit consisted of the 16S ribosomal RNA and 22 ribosomal proteins (Table S1).

1.1.2 *Modeling of the missing residues of proteins*

The terminal residues of proteins have often not been determined experimentally. Although the absence of some of the residues could not significantly affect the results of the simulations, the C-terminus of uS9, uS13, and uS19 was considered to interact with the initiator tRNA.

Table S1. Molecules that are contained in the preinitiation complex.

| | |
|---|---|
| RNA | 16S ribosomal RNA<br>initiator tRNA<br>mRNA |
| Ribosomal proteins | uS2, uS3, uS4, uS5, uS7, uS8, uS9, uS10,<br>uS11, uS12, uS13, uS14, uS15, uS17, uS19<br>bS1, bS6, bS16, bS18, bS20, bS21, bTHX |
| Initiation factors | IF1[a], IF3 |

[a]IF1 was not included in the simulation system of PIC-4 because it was not included in the experimental structure of PIC-4 (pdb id: 5lmu).

To model amino acid residues not determined by the experiments, we performed structure modeling using MODELLER [2] and Chimera [3]. We used "Model loops" in Chimera and the DOPE score to select a model structure. In PIC-2A, we generated ten candidate structures of the C-terminal region of uS19. Since some of these generated structures overlapped with other molecules in the PIC, we excluded such candidates. Finally, we selected the structure with the best score from the candidate structures.

The structure modeling protocols for PIC-3 and PIC-4 were almost the same as for PIC-2A. Because the C-terminal region of uS13 was also missing, we performed structure modeling not only for uS19 but also for uS13. In addition, there were four internal missing residues in IF3. We also performed loop modeling for the segment in IF3 and selected the loop structure with the best score from the generated structures.

##### 1.1.3 *Preparation of tRNA molecules*

Some tRNAs had a modified base in the experimental structures. Because these RNAs cannot be treated in the Martini model, we replaced the base of the tRNAs with a standard base. RNA residues PSU, 4SU, 5MU, OMC, and G7M were replaced by U, U, U, C, and G, respectively.

#### 1.2 Coarse-grained model for the simulation system

##### 1.2.1 *Coarse-grained model of the PIC using the Martini force field*

We prepared the all-atom model of the PIC as described above. Then we converted the all-atom model into a coarse-grained model. We used the Martini 2.2 force field for proteins [4-6], RNAs [7], ions, and solvent. In the Martini model, several atoms are mapped to one coarse-grained bead. This mapping was done using the script martinize.py. The resulting coarse-grained PIC using the Martini model consisted of approximately 16,000 beads.

The PIC system contained zinc and magnesium ions. These ions are not defined in the Martini model. Therefore, we added the definition of zinc and magnesium ions to the parameter file of the Martini force field. We used bead type Qd as their bead type and set the electric charge of the ions to +2.

Coarse-grained beads corresponding to water and ions were then added to the coarse-grained PIC system. The PIC system was placed in a rhombic dodecahedron box. The dimensions of the simulation box were set so that the distance between the PIC and the box boundary was at least 4.0 nm. Periodic boundary conditions were applied. Coarse-grained beads corresponding to water were added to the box. Sodium and chloride ions were also added to the simulation box so that the ion concentration was 0.15 M and the

Table S2. Simulation systems of the PIC.

| | PIC-2A (5lmq) | PIC-3 (5lmt) | PIC-4 (5lmu) |
|---|---|---|---|
| Number of residues | | | |
| 16S rRNA | 1,514 | 1,515 | 1,515 |
| mRNA | 21 | 20 | 20 |
| tRNA | 77 | 77 | 77 |
| Proteins | 2,644 | 2,642 | 2,571 |
| Number of beads | | | |
| RNAs | 10,579 | 10,579 | 10,579 |
| Proteins | 5,873 | 5,868 | 5,710 |
| Water | 161,125 | 152,266 | 151,436 |
| $Na^+$ | 3,177 | 3,042 | 3,040 |
| $Cl^-$ | 1,968 | 1,876 | 1,858 |
| $Zn^{2+}$ | 2 | 2 | 2 |
| $Mg^{2+}$ | 64 | 86 | 80 |
| Total | 182,788 | 173,719 | 172,705 |

simulation system was electrically neutral. Finally, the total number of beads in the simulation system was approximately 170,000. Table S2 summarizes the simulation systems of the PIC.

1.2.2 *Elastic network model for proteins and RNAs*

We added restraints between the intramolecular beads of proteins and RNAs using an elastic network model [8] because some of the energy minimization calculations and equilibrium simulations were unstable without restraint potentials. We used a harmonic potential as the restraint potential. The restraint potentials were imposed to restrict the distance between the beads to the distance in the initial structure.

For proteins, we restrained coarse-grained beads corresponding to the backbone atoms of the protein if the distance between the beads was less than 0.8 nm. We used the initial position of the beads to calculate the distances. The force constant of the harmonic potential was set to 500 kJ mol$^{-1}$ nm$^{-2}$. For the C-terminal region of uS19, we excluded the restraint potentials because the position of these beads was determined by the modeled structure and the C-terminal region was considered to be flexible.

For RNAs, we restrained coarse-grained beads if the distance between the beads was less than 1.0 nm. The initial position of the beads was used to calculate the distances. The force constant of the harmonic potential was set to 500 kJ mol$^{-1}$ nm$^{-2}$. Because the beads that satisfy the distance criteria were restrained, this restraint and other intramolecular interactions, such as the bond and angle energy terms, interfered with each other. Therefore, we used only the elastic network model as the intramolecular interactions of RNAs. In this study, because we focused on analyzing the intermolecular interactions of the PIC, this interaction model of RNAs was considered valid.

### 1.3 Protocols for the MD simulations

#### 1.3.1 *Energy and force calculations*

Potential energy and force were evaluated using the Martini force field. The intramolecular interactions of RNAs were excluded except for the energy and force of the elastic network model (see 1.2.2). The cutoff distance for non-bonded interactions was set to 1.1 nm. The electrostatic interactions were calculated with reaction field [9]. The parameters of reaction field $\varepsilon_f$ and $\varepsilon_{rf}$ were set to 15.0 and 0.0, respectively.

#### 1.3.2 *Energy minimization*

We performed energy minimization for all PIC systems using GROMACS 2020.4 [10]. A steepest descent algorithm was used for energy minimization. The minimization tolerance was set to 100 kJ mol$^{-1}$ nm$^{-1}$. The maximum number of minimization steps was set to 5,000. During the energy minimization, the backbone beads of the proteins and all RNA beads were restrained to the initial structure using a harmonic potential with a force constant of 1,000 kJ mol$^{-1}$ nm$^{-2}$.

#### 1.3.3 *MD simulations for equilibration*

After minimizing the energy of the coarse-grained PIC systems, we performed MD simulations for equilibration using GROMACS 2020.4 [10]. The reference temperature and pressure were set to 310 K and 0.1 MPa, respectively. The temperature was controlled by the velocity rescaling algorithm with a stochastic term [11]. We set the time constant for the temperature control to 2.0 ps. The pressure was controlled by the weak coupling algorithm with isotropic scaling [12]. We set the time constant and compressibility for the pressure control to 10.0 ps and 0.003 $MPa^{-1}$, respectively. The initial velocity of each bead was generated using the Maxwell-Boltzmann distribution at 310 K.

The MD simulations for equilibration consisted of seven steps.

(1) MD simulations were performed for 1.0 ns. The time step for integration was set to 10 fs. The backbone beads of the proteins and all RNA beads were restrained to the minimized structure using a harmonic potential with a force constant of 1,000 kJ $mol^{-1}$ $nm^{-2}$.

(2) MD simulations were performed for 20.0 ns. The time step for integration was changed to 20 fs. The time step of subsequent steps was set to the same value. The backbone beads of the proteins and all RNA beads were restrained to the structure obtained in (1) using a harmonic potential with a force constant of 1,000 kJ $mol^{-1}$ $nm^{-2}$.

(3) MD simulations were performed for 20.0 ns. The backbone beads of the proteins and all RNA beads were restrained to the structure obtained in (2) using a harmonic potential with a force constant of 200 kJ $mol^{-1}$ $nm^{-2}$.

(4) MD simulations were performed for 20.0 ns. The backbone beads of the proteins and all RNA beads were restrained to the structure obtained in (3) using a harmonic potential with a force constant of 40 kJ $mol^{-1}$ $nm^{-2}$.

(5) MD simulations were performed for 20.0 ns. The backbone beads of the proteins and all RNA beads were restrained to the structure obtained in (4) using a harmonic potential with a force constant of 8 kJ $mol^{-1}$ $nm^{-2}$.

(6) MD simulations were performed for 20.0 ns without the restraints.

(7) MD simulations were performed for 20.0 ns without pressure control (i.e., with constant volume).

### 1.4 Metadynamics and umbrella sampling simulations

#### 1.4.1 *Reaction coordinates*

We defined the distance between the tRNA and the small ribosomal subunit as the reaction coordinate (DIST). This reaction coordinate was used for both metadynamics [13] and umbrella sampling [14]. This reaction coordinate characterizes the dissociation process of the initiator tRNA from the ribosome. First, we defined the set of tRNA beads that were within 0.8 nm of the ribosomal RNA in the initial structure (Group 1). Similarly, we defined the set of ribosomal RNA beads that were within 0.8 nm of the initiator tRNA in the initial structure (Group 2). We then calculated the center of mass for the two groups and defined the reaction coordinate as the distance between the two centers.

We then defined another reaction coordinate (CN) using a coordination number. This reaction coordinate was used for metadynamics only. The coordination number was obtained by calculating the number of contacts between Group 1 and Group 2. The number of contacts between the two groups was calculated using a switching function with parameter $R_0$ equal to 2.0 nm [15,16].

#### 1.4.2 *Metadynamics simulations*

We performed metadynamics simulations [13] for all PIC systems using the structure obtained by the equilibration. These metadynamics simulations were performed using GROMACS 2020.4 [10] and PLUMED 2.6.2 [15,16]. The temperature of the system was controlled and set to 310 K. The volume of the system was fixed. The time step of integration was set to 20 fs. Each metadynamics simulation was performed for 500 ns. For each simulation, the initial velocity of each bead was generated using the Maxwell-Boltzmann distribution at 310 K.

We used the two reaction coordinates (DIST and CN) defined above (see 1.4.1) for the metadynamics simulations. Therefore, we performed two-dimensional metadynamics simulations. The first reaction coordinate DIST was used to estimate the tRNA dissociation path from the ribosome, while the second reaction coordinate CN was used to enhance the structure sampling. The width of the Gaussian hills for DIST and CN was set to 0.05 nm and 10.0, respectively. The height of the Gaussian hills was set to 0.5 kJ mol$^{-1}$. The frequency for the Gaussian hill addition was set to 0.1 ps$^{-1}$. We also added a restraining harmonic potential that starts acting on the system when the reaction coordinate DIST is larger than 3.0 nm, so that the tRNA does not go far from the ribosome. The force constant of the potential was set to 1,000 kJ mol$^{-1}$ nm$^{-2}$. The number of the metadynamics simulations for PIC-2A, PIC-3, and PIC-4 was 6, 8, and 6, respectively. The number of trajectories for PIC-3 was larger than that for the others because the convergence for PIC-3 was not as good as that for PIC-2A and PIC-4 (see the error bars of the free energy in Fig. 2c in the main text).

From the metadynamics trajectories, we extracted structures every 0.05 nm in the range of 0.5 nm (0.6 nm for PIC-4) to 1.5 nm for the reaction coordinate DIST. For example, when we found the simulation time when the value of DIST was 1.0 nm for the first time, we extracted the structure at that time. We treated this structure as the one when the reaction coordinate DIST was equal to 1.0 nm. These structures were used for the initial structure of the umbrella sampling simulations.

#### 1.4.3 *Umbrella sampling simulations*

We performed umbrella sampling simulations [14] for all PIC systems using the structures from the metadynamics trajectory. These umbrella sampling MD simulations were performed using GROMACS 2020.4 [10] and PLUMED 2.6.2 [15,16]. The temperature of the system was controlled and set to 310 K. The volume of the system was fixed. The time step of integration was set to 20 fs. Each umbrella sampling simulation was performed for 500 ns. For each simulation, the initial velocity of each bead was generated using the Maxwell-Boltzmann distribution at 310 K.

We used DIST as the reaction coordinate of the umbrella sampling simulations. We used a harmonic potential with a force constant of 5,000 kJ $mol^{-1}$ $nm^{-2}$ as the umbrella potential of the simulations. The umbrella sampling simulations were performed with the reference distance of the harmonic potential set to every 0.05 nm in the range of 0.5 nm (0.6 nm for PIC-4) to 1.5 nm along the tRNA dissociation path from the metadynamics simulation. As a result, the number of the umbrella sampling simulations using one metadynamics trajectory was 21, 21, and 19 for PIC-2A, PIC-3, and PIC-4, respectively. Since umbrella sampling simulations along the tRNA dissociation path were performed for all the metadynamics trajectories, the total simulation time for PIC-2A, PIC-3, and PIC-4 was 63.0 μs (21 × 6 × 0.5 μs), 84.0 μs (21 × 8 × 0.5 μs), and 57.0 μs (19 × 6 × 0.5 μs), respectively.

#### 1.5 MD simulations without electric charges of the C-terminal region of uS19

We performed MD simulations for PIC-2A and PIC-4 in which the electrostatic interactions caused by the C-terminal region of uS19 were eliminated. We used the same system preparation and coarse graining protocols as described in 1.1 and 1.2. To eliminate the electrostatic interactions in the C-terminal region, the electric charge of the beads of the last 10 amino acid residues (GKEAKATKKK) in the C-terminal region was modified to zero in the parameter file of uS19 (see Fig. 4a in the main text). Because the net electric charges of the system became negative due to the charge elimination, the chloride ions were eliminated to neutralize the system. The number of chloride ions for the system of PIC-2A and PIC-4 was 1,965 and 1,855, respectively. For these systems, energy minimization and MD simulations for equilibration were performed using the same procedures as described in 1.3.

Metadynamics and umbrella sampling simulations were performed to calculate the dissociation free energy of the initiator tRNA. We also used the same simulation conditions as described in 1.4. The number of the metadynamics simulations for PIC-2A and PIC-4 was 6 and 7, respectively. The number of the umbrella sampling simulations using one metadynamics trajectory was 21 for both PIC-2A and PIC-4. As a result, the total simulation time for PIC-2A and PIC-4 was 63.0 μs (21 × 6 × 0.5 μs) and 73.5 μs (21 × 7 × 0.5 μs), respectively.

### 2. Additional Results for the Molecular Dynamics Simulations

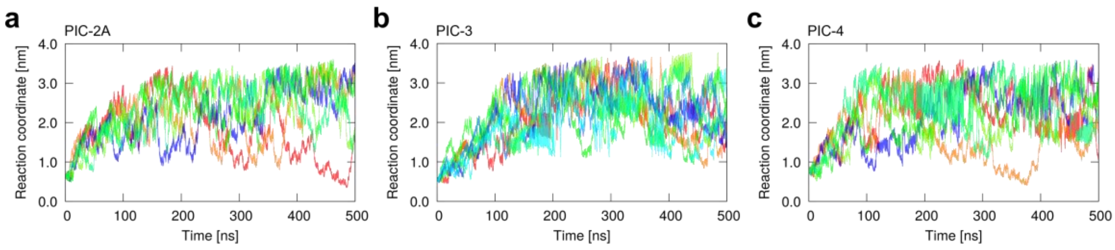


Figure S1. Results of the metadynamics simulations for PIC-2A, PIC-3, and PIC-4. The time series of the reaction coordinate DIST for the metadynamics simulations in (a) PIC-2A, (b) PIC-3, and (c) PIC-4 are shown. The different time series are displayed in different colors.

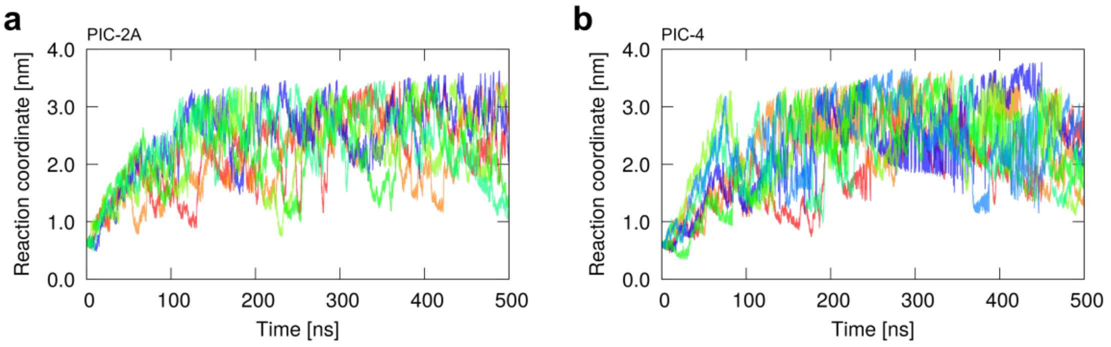


Figure S2. Results of the metadynamics simulations in which the electric charges of the C-terminal residues of uS19 were set to zero. The time series of the reaction coordinate DIST for the metadynamics simulations in (a) PIC-2A and (b) PIC-4 are shown. The different time series are displayed in different colors.

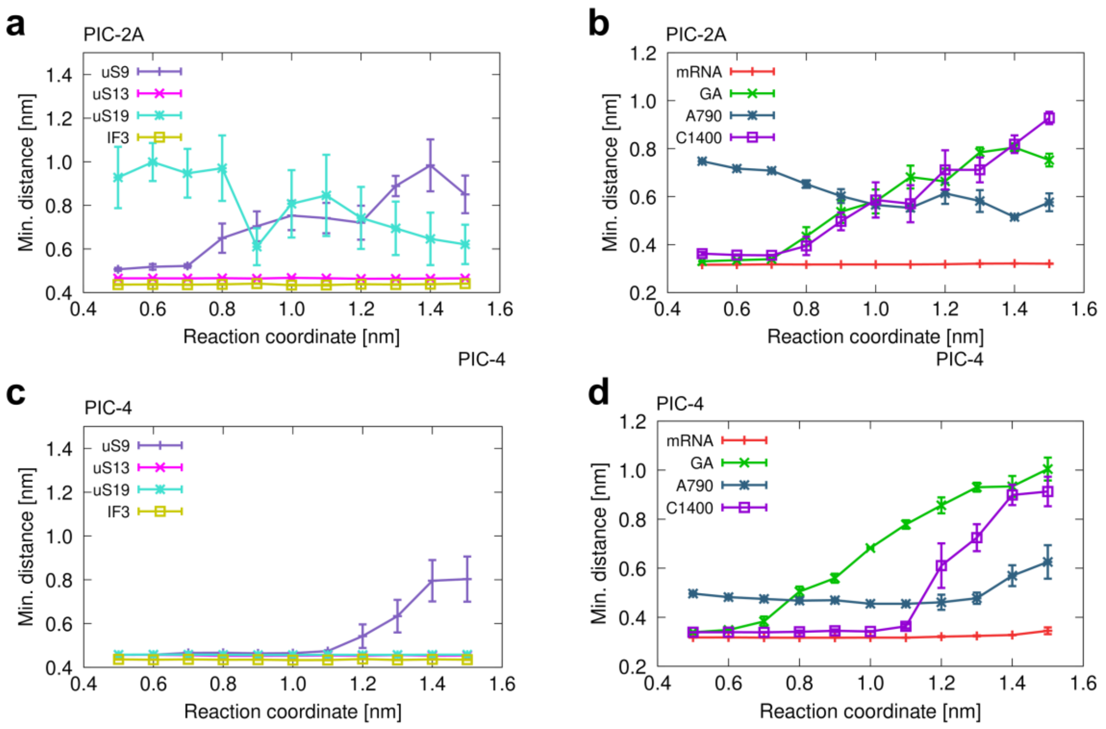


Figure S3. Results of the metadynamics simulations in which the electric charges of the C-terminal residues of uS19 were set to zero. The distances between the tRNA and the ribosomal proteins are shown as a function of the reaction coordinate DIST for (a) PIC-2A and (c) PIC-4. The distances between the tRNA and the mRNA and ribosomal RNA residues are shown as a function of the reaction coordinate DIST for (b) PIC-2A and (d) PIC-4. The error bars represent the standard error of the distance.

[11] Bussi, G.; Donadio, D.; Parrinello, M. Canonical Sampling through Velocity

Rescaling. *J. Chem. Phys.* **2007**, *126* (1), 014101.
